# How much storage precision can be lost: Guidance for near-lossless compression of untargeted metabolomics mass spectrometry data

**DOI:** 10.1101/2023.03.14.532504

**Authors:** Junjie Tong, Miaoshan Lu, Bichen Peng, Shaowei An, Jinyin Wang, Changbin Yu

## Abstract

The size of high-resolution mass spectrometry (HRMS) data has been increasing significantly. Several lossy compressors have been developed for higher compression rates. Currently, a comprehensive evaluation of what and how MS data (*m/z* and intensities) with precision losses would affect data processing (i.e., feature detection and compound identification) is absent.

Here, we set an error threshold at 1% to assess the significance of the difference between two files in feature and compound detection results obtained from MZmine3. First, we examined that mzML files with both *m/z* and intensity encoded in 32-bit precision appear to be a preferred combination via msConvert, which has smaller file size and minor variation with other combinations of storage precision (<0.13%). We then identified that the absolute error of 10^−4^ for *m/z* had a feature detection error of 0.57% and compound detection error of 1.1%. For intensities, the relative error group of 2×10^−2^ had an error of 4.65% for features and 0.98% for compounds, compared with precision-lossless files. Taken together, we provided a reasonable scene-accuracy proposal, with a maximum absolute error of 10^−4^ for *m/z* and a maximum relative error of 2×10^−2^ for intensity. This guidance aimed to help researchers in improving lossy compression algorithms and minimizing the negative effects of precision losses on downstream data processing.

## 1. Introduction

Untargeted metabolomics, as a study of measuring the metabolome within an organism, has been extensively applied for disclosing novel biomarkers clinically. In this field of study, liquid chromatography-mass spectrometry (LC-MS) technology is considered the prime choice for small molecule identification and quantification. To exploit the information from a raw MS output file requires a workflow of data analysis, including data processing, data annotation, statistical analysis, functional analysis, etc., [1]. Among them, data processing is a critical step and the cornerstone of untargeted metabolomics data analysis, which comprises extracted ion chromatogram (EIC), peak detection, peak deconvolution, and peak alignment [1]. Several bioinformatics tools and platforms have been introduced to address the needs of processing untargeted metabolomics data, some of which are open-source software, like MZmine3 [2], MS-Dial [3], XCMS [4], MetaboAnalyst 5.0 [5], and other commercial tools, such as Compound Discoverer (Thermo Scientific), MarkerView (SCIEX), Mass Profiler Professional (Agilent), MarkerLynx (Waters).

Raw vendor files from the mass spectrometer, for example, the .Raw extension for Thermo and .wiff extension for SCIEX instruments, are lacking cross-platform compatibility and software flexibility [6]. As a result, a community standard format, “mzML”, has been developed and rapidly become the most commonly open format for MS data [7]. One of the disadvantages of mzML is the much bigger file size than vendor format, resulting in a higher cost of storage. Thus, a number of compression algorithms for MS data have been proposed recently. In general, these compression strategies could be classified into lossless, such as MassComp [8], and lossy (or “near-lossless”), like MSNumpress [9], Aird-ZDPD [10], or both, like mspack [11]. These lossy algorithms achieved higher compression rates at the expense of certain storage precision of raw data. The following question is how much precision losses of raw data (i.e., *m/z* and intensities) will affect the downstream data processing to some acceptable or unacceptable extent. In MSNumpress, Teleman et al. reported a maximum relative error of 2×10^−9^ (0.002ppm) for *m/z* and 2×10^−4^ for intensities generated the identical peptide list (FDR < 1%) as the raw files. For mspack, Hanau et al. showed that a compressed file with a maximum absolute error of 10^−4^ (0.2ppm at 500 *m/z*) for *m/z* and a relative error of 10^−2^ for intensities would be well below the mass accuracy of mass analyzers and the difference between repetitions of an experiment, respectively. In a recent study, Wang et al. developed Stack-ZDPD from ZDPD and observed that the *m/z* data stored at 5 decimal places (0.1ppm at 100 *m/z*) generate the almost same peak lists detected via MZmine3 [12]. Till now, there is no study has systematically evaluated the impacts of data precision losses of *m/z* and intensities data in mzML files on subsequent data processing step, specifically feature detection and compound annotation, with the former describing the processing results of the original signal while the latter is more dependent on the scope of the search library.

In this study, we first converted metabolomics MS vendor files into precision-lossless mzML files via command line tools of msConvert [13], where *m/z* values and intensities were encoded in four combinations of storage precision (32- or 64-bit). Then, we tested the differences of these combinations based on the critical results of feature and compound identification. We used a two-dimensional window of RT shift <= 0.025min and *m/z* error <= 5ppm to determine whether two features are identical. Furthermore, we generated precision-lossy mzML files with a defined maximum absolute error of *m/z* or a relative error of intensities by the truncation transformations mentioned in mspack. These truncated mzML files were also tested in terms of feature and compound detection. Eventually, we compared precision-lossy files to precision-lossless files in the total number of features, consensus features ratio, intensity fold change (FC) of consensus features, and annotated consensus compounds.

Considering the mass accuracy of current instruments and reproducibility of an experiment, we set the threshold value of the error to 1%. After comparing the above differences with the error threshold, we proposed a guidance for a suitable storage precision combination in precision-lossless mzML files, and for the storage precision losses of *m/z* and intensity value in precision-lossy compression.

## 2. Experimental Section

### 2.1 Vendor Files Preparation

In order to complete a comprehensive evaluation, we selected sixteen LC-MS vendor files (labeled from Met_01 to Met_16) generated by different types of instruments, from various organisms, and acquired by different acquisition methods (Table s1). All vendor files can be searched on the open-access database repository of Metabolights (https://www.ebi.ac.uk/metabolights/).

To determine the suitability of files, we required them must meet the inclusion criteria. (1), It must be the raw vendor file rather than the derived file format (e.g., mzML, mzXML), because the storage precision of mzML files has been fixed. (2), These files were generated from untargeted metabolomics or lipidomics studies as the workflow of MZmine3 is designed for untargeted metabolomics data processing. (3), The MS data from high-resolution mass spectrometers is required as the mass detection algorithm is suitable for HRMS data.

### 2.2 Precision-lossless Files Encoded in 32-bit and 64-bit precision

Sixteen vendor files were converted to the open format, mzML, by the command line tool of msConvert (ProteoWizard) [13]. Each vendor file generated four mzML files with *m/z* and intensities encoded in different combinations of storage precision: (1), both *m/z* and intensity values were stored at single (32-bit) precision, with parameters “--mz32 --inten32” in msConvert. (2), both *m/z* and intensity data with double (64-bit) precision, parameters as “--mz64 --inten64”. (3), *m/z* and intensities encoded in 32-bit and 64-bit precision, respectively, parameters as “--mz32 --inten64”. (4), *m/z* at 64-bit precision and intensities at 32-bit precision (“--mz64 --inten32”).

### 2.3 Truncation Transformations for m/z and Intensity Values

The truncation transformations idea is one of the lossy compression methods used in mspack compressors [11]. According to the IEEE 754 standard format of a floating-point number (Fig. 1), the single-precision floating numbers contained one sign bit, eight exponent bits, and 23 fraction bits, for example, a binary format of 32-bit floating point number could be expressed by (−1)^*sign*^2^*exponent*^x1.*fraction*. The idea of truncation transformation is to set the last significant bits in the fraction part to zero, which creates a certain degree of precision losses on the import value.

**Fig. 1.**
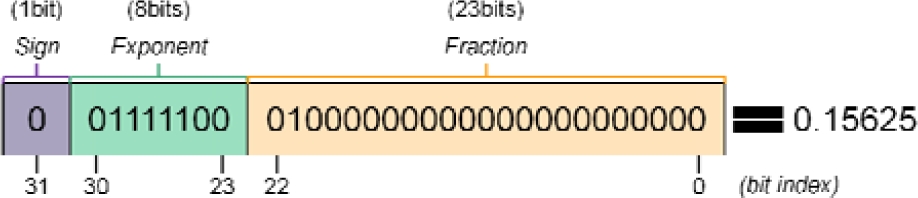
This figure is adopted from [13], which shows the binary format of value 0.15625 in IEEE 754 standard with single precision. The value can be computed from the binary representation as 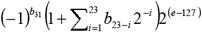, with 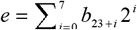.

In the MS data, *m/z* and intensity values could be transformed to data with defined errors by this idea. We transformed these precision-lossless files from Chapter 2.2 and created lossy files with a certain degree of maximum absolute error of *m/z* or relative error of intensity data. The precision-lossy files were used for the downstream data processing workflow. This homemade truncation transform code, named PrecisionControlTask, was written in Java language and was locally integrated into the IO module (import_rawdata_mzml) of a forked Java-based MZmine3 source code (https://github.com/TongJJCSi/mzmine3).

#### 2.3.1 The truncation transformations for *m/z* values (absolute errors)

The input *m/z* floating-point values (32-bits or 64-bits) were converted to their binary format under IEEE754 standard. Based on the defined absolute error, we calculated the fraction bits needed to be truncated (details of bits calculation can be found here [11]) and set them to zero. Subsequently, each precision-lossless mzML file was transformed to five precision-lossy mzML files with different degrees of absolute errors in *m/z*, resulting in maximum absolute errors of 10^−5^, 2×10^−5^, 5×10^−5^, 10^−4^ and 10^−3^. The sixteen mzML files with the same absolute errors were regarded as an error group. These error groups were selected with reference to previous studies, which showed that MSNumpress (2×10^−9^ or 0.002ppm) and mspack (10^−4^ or 0.2ppm at 500 *m/z*) defined the maximum relative and absolute errors for *m/z* without negative effects on the following data processing. Therefore, we first chose an intermediate absolute error of 10^−5^ (0.02ppm at 500 *m/z*) that was examined no difference with precision-lossless files (error threshold < 1%), and then extended error range to 10^−3^, where significant difference between precision-lossless and -lossy files were occurred.

#### 2.3.2 The truncation transformations for intensity values (relative errors)

Similar to the truncation transformation used for *m/z* values. In order to control the maximum relative error of intensities, the way of calculating the number of truncated bits was referenced to the idea in mspack [11]. Then, we set five error groups with a range of maximum relative errors, 2×10^−4^, 2×10^−3^, 8×10^−3^, 2×10^−2^ and 2×10^−1^. All files with the same relative errors were clustered into an error group. The reason for setting these groups is similar to *m/z* Based on the maximum relative errors in MSNumpress (2×10^−4^) and mspack (10^−2^), we checked the range of relative errors from 2×10^−4^ to 2×10^−1^, describing the trend of variation in feature and compound identification is getting bigger as the expansion of maximum relative errors of intensity data.

In general, each precision-lossless file was transformed into ten lossy files (five with only *m/z* errors and five with only intensity errors)

### 2.4 LC-MS Data Processing Workflow via MZmine3

MZmine 3 (v3.2.8) was used to process both precision-lossless (files from Chapter 2.2) and precision-lossy (files from Chapter 2.3) mzML files. All parameters used in MZmine3 can be seen in Table s2 (Supplementary). All mzML files were imported into MZmine3 in profile mode (except Met-11 in centroided mode), so the first step was the mass detection module for data-centroiding. In this module, the algorithm of exact mass was used to centroid the profile data. In order to obtain more peaks, we set the noise level at 1. The noise level set a noise threshold and excluded background noise under the defined level, which is dependent on the type of mass spectrometers. Thus, the recommended way of setting noise level is reviewing each file, but it may still miss some signals. Therefore, we set the value at 1 to include more features and set another noise-filter downstream.

Afterwards, the EIC building was processed by the Automated Data Analysis Pipeline (ADAP) chromatogram builder, where the scan-to-scan accuracy (*m/z*) parameter was set to 10ppm for orbitrap LC-MS data and 15ppm for time-of-flight (TOF) generated files. Then, peaks in these EICs were detected by ADAP feature resolver module with default parameters, where Signal-to-Noise was estimated in an intensity window-based method. The feature alignment was performed in a default setting as well.

After all, each feature list was searched against an online database via the integrated annotation module in MZmine3. The determination of the database was based on the organisms’ type of mzML files, such as the human samples were against Human Metabolome Database (HMDB) (Met_01, 02, 08, 09, 10, 11, 12, 13, 14), the yeast sample against Yeast Metabolome Database (YMDB) (Met_15), and the remaining samples against Kyoto Encyclopedia of Genes and Genomes (KEGG) metabolites database (Met_03, 04, 05, 06, 07, 16). The feature lists annotated by the metabolites were exported as .csv format and stored locally for further data statistics.

### 2.5 Data Statistics of Precision-lossless Files

The feature list and annotated compound list of each precision-lossless file (from Chapter 2.2) were exported via MZmine3. For feature lists, we defined a feature via its *m/z* and retention time (RT) value. Then, we examined the overlapping features (consensus features) of four files and presented them as Venn diagrams, which are generated from the same vendor file but with different combinations of storage precision. Next, we calculated the proportion of unique features for each file to demonstrate the difference between these combinations. To avoid mistakenly assigning “real” consensus features with a minor margin of error as a unique feature, we applied a window of *m/z* <= 5ppm and RT shift <= 0.025min to filter consensus features. After the filtration, we recalculated the filtered fraction of unique features for each file.

Downstream of feature detection, we defined a compound through its compound name, molecular formular, and neutral mass information from compound lists. In each file, we counted the total number of annotated compounds and screened out duplicate compounds. Afterward, we calculated the unique compound rate (UCR) in all files.

After all, we evaluated the differences in feature and compound detection between storage precision combinations via the threshold value of error (1%).

### 2.6 Data Statistics of Truncated Files

All *m/z* and intensity precision-truncated files (from Chapter 2.3) were processed via MZmine3 and generated feature and compound lists. The feature and compound detection results of truncated files were parallelly compared to the results of precision-lossless files (from Chapter 2.5). The overlapping features or compounds between precision-lossy and -lossless files were regarded as consensus features or compounds while these only detected in precision-lossy files are called unique features or compounds.

In feature lists, the number of total features and consensus features were calculated. Then, we checked the filtration results of consensus features under a window of *m/z* <= 5ppm and RT shift <= 0.025min. After filtration, we calculated the ratio of unique features to total features of precision-lossless files, which was compared to the acceptable margin of error (1%). Furthermore, we measured the distribution of fold-change (FC) of intensity in consensus features. In compound lists, the total number and percentage of consensus compounds across all files were calculated to evaluate the impacts of different degrees of precision losses on compound detection.

## 3. Results and Discussion

### 3.1 Single- Vs Double-Precision Storage for m/z and Intensities in mzML Files

Converting raw vendor files to mzML files using msConvert with default settings would encode the *m/z* values in 64-bit precision (i.e., double-precision) and intensity values in 32-bit precision (i.e., single-precision) for the output files [12]. However, in some data processing software, both *m/z* and intensity data are calculated in double precision. Therefore, we aim to test and evaluate the impact of the degree of variation of mzML files at four different combinations of storage precision by comparing their feature and compounds identification results exported from MZmine3.

#### 3.1.1 Comparison of Feature Detection Results

A feature is defined as a two-dimensional signal (i.e., *m/z* and retention time) [14]. Thus, we recognized a non-zero feature with identical *m/z* and retention time (RT) overlapping across multiple mzML files to be a consensus feature. We observed that files have the identical feature list regardless of storage precision of intensity values when *m/z* values are encoded in a fixed precision (Supplementary Fig. s1). However, the storage precision of intensity data at a fixed point, *m/z* values would generate a feature list with differences between 32-bit and 64-bit precision. We calculated the unique feature rate (UFR), the percentage of unique features to total features, to evaluate the degree of difference between files with *m/z* at 32- or 64-bits precision. Under the definition of consensus features (*m/z* and RT values are identical in two files), the average UFR between files with fixed precision of intensity and unfixed precision of *m/z* values is about 3.3% (see unfilter lines in Fig. 2a). This unfiltered UFR showed a noticeable degree of difference (>1% of threshold value of error) in feature lists of files.

**Fig. 2.**
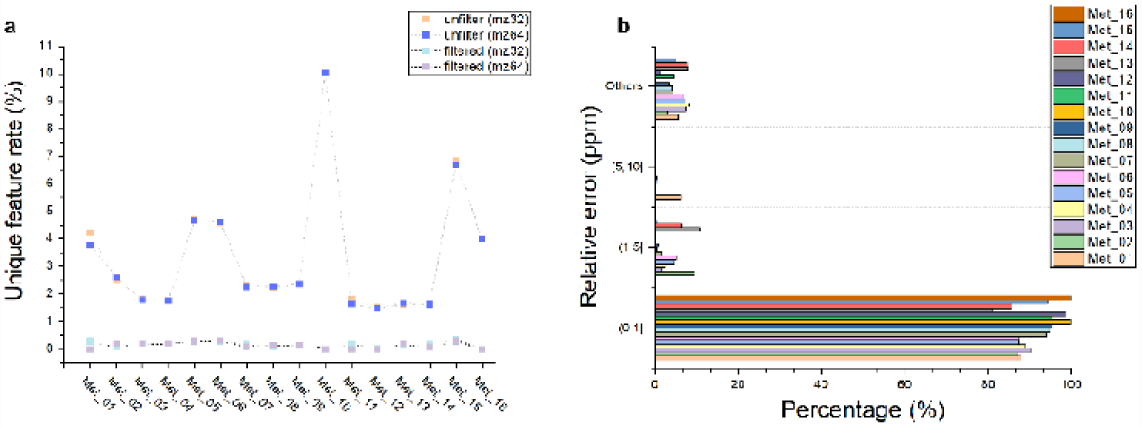
The filtration process of unique features in both 32-bit and 64-bit precision mzML files. (**a**) The unique feature rate in all files before and after filtration; (**b**) The distribution of relative errors of unique features in *m/z* 32-bit and 64-bit replicates of all 16 files within a RT shift <= 0.025min. The group of “others” comprises features with RT shift > 0.025min or *m/z* error > 10 ppm.

However, we noticed that a part of unique features with a minor margin of difference to consensus features. A feature occurs across repetitions of an experiment may bring a RT drift (few seconds to tens of seconds) problem, or *m/z* errors due to mass analyzers. Some studies suggested that RT shift and *m/z* errors could be corrected via a certain two-dimensional window of *m/z* and RT, such as the window of RT shift < 1.5s (0.025min) and *m/z* error < 0.01 (20ppm at 500 *m/z*) [15]. We set the RT shift less than 1.5s (0.025min), where the distribution of unique features, from *m/z* 32- and 64-bit precision files, over different relative error (ppm) ranges was examined (Fig. 2b). We identified that over 90% of unique features in all files could be reassigned to consensus features with a window of *m/z* error <= 5 ppm and continuing to expand this window to 10ppm has a limited help in reassigning more consensus features. Therefore, we used a window of RT <= 0.025min and *m/z* error <= 5 ppm to screen all unique features. Consequently, the filtered UFR were adjusted to a negligible level across all files, with an average rate of 0.13% (see filtered lines in Fig. 2a).

However, the discrepancy needed to be further evaluated on the compound annotation list to determine if the difference in feature lists carries over to the list of identified compounds. As files with fixed storage precision of *m/z* and 32-bit or 64-bit precision of intensities generate the same feature lists, we then evaluated differences between files with *m/z* values of 32-bit and 64-bit precision in the following compound identification.

#### 3.1.2 Comparison of Online Compound Database Search Results

Feature lists of all files were processed via the online compound database search module of MZmine3. The total number of detected compounds ranges from 3 to 9765 among sixteen files (Table s3). A compound is defined by its compound name, molecular formula, and neutral mass. After screening out these repeated compounds, the number of compounds saw a decrease. Next, we calculated the UCR for all files, except for Met_10, Met_15, and Met_16 files with a compound number less than 50, resulting in an average UCR of 0.13%.

Overall, we considered that the variation on feature (0.13%) and compound (0.13%) detection across different combinations of *m/z* and intensity storage precision is well below the error threshold (1%). Thus, the files with both *m/z* and intensity values encoded in 32-bit precision were regarded as a preferred choice when researchers require smaller file sizes and faster data analysis. In the following study, we selected these files (mz32intensity32) for truncation transformation to examine a suitable precision loss for *m/z* and intensity value.

### 3.2 Truncation Transform of Intensity Values

We aimed to test the changes of feature and compound lists as the relative errors of intensities increase. Here, we set five groups (2×10^−4^, 2×10^−3^, 8×10^−3^, 2×10^−2^, 2×10^−1^) by truncation transformation to identify the point of relative error with or without changes for data processing.

#### 3.2.1 Comparison of Feature Lists among Intensity Groups with Different Relative Errors

To assess the differences between groups, we set the precision-lossless files as the “Blank” group, Features shared between precision-lossless and lossy files were considered as “Consensus features”, features detected only in precision-lossy files were regarded as “Unique features”.

For unfiltered files, the number of total features had a significant decrease occurred in the group of all files with a relative error of 2×10^−1^ (p < 0.05) (Table S3). For consensus features, a noticeable decrease started in the group of 8×10^−3^ (Fig. s2). Afterwards, we calculated the ratio of consensus features to total features in the blank file (except for Met_10 has only 10 features) under the window of RT <= 0.025min and *m/z* error <= 5 ppm (Fig. 3). With the filtration window, the proportion of consensus features reached over 95% in all groups (2×10^−4^ for 99.96%, 2×10^−3^ for 99.63%, 8×10^−3^ for 98.45%, 2×10^−2^ for 95.35%), except for 2×10^−1^ (62.78%). For the filtered consensus features, we then calculated the fold-change (FC) of their intensities between blank files and error groups, which may impact the compound identification in the downstream of data analysis (Fig. s3).

**Fig. 3.**
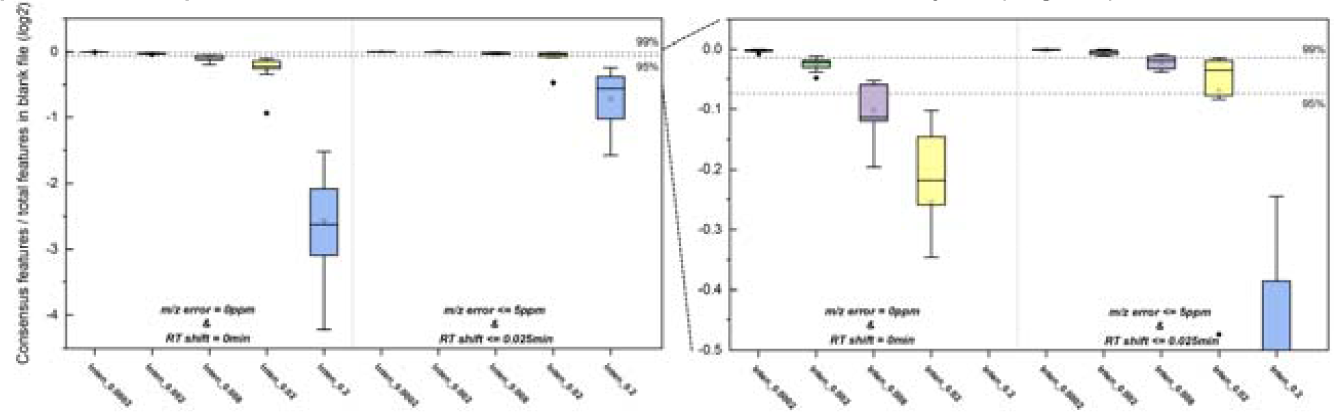
The ratio of consensus features to total features of blank files in each group with truncated intensities under different filtration windows. (*m/z* & RT error equals 0, *m/z* <= 5ppm & RT <= 0.025min). The ratio values were *log*_*2*_ transformed. Met_10 was excluded due to the number of total features (n=10) was too small.

In general, the error group of 2×10^−1^ could substantially change the feature list of the blank file. The proportion of unique features is less than 1% of error threshold in two groups (2×10^−4^, 2×10^−3^), which achieved a good performance when the original signal was processed for identification of features. Moreover, the fraction of significant consensus features (FC>2 or FC<0.5) was considered in a controllable level in all groups (less than 1%).

In MS data processing studies, chromatogram construction of EIC and chromatographic peak detection are critical steps and are prone to generate false positive or false negative features [15]. Tracing back to modules used in our study, three of them (i.e., Mass detection, ADAP chromatogram builder, ADAP feature resolver) are likely related to the discovery of unique features. In the mass detection module, due to these files being in profile mode, the algorithm of Exact mass is selected for data centroiding. The errors of intensities might shift the location of data points closest to the peak center at half maximum intensity, which causes some errors in exact mass detection. For the module of ADAP chromatogram builder, two user-defined parameters (group intensity threshold and min highest intensity) might include some false EICs or exclude true EICs due to the changes of data points intensity values. In the ADAP peak detection module, we set the parameter of the Signal/Noise estimator to an intensity-based window method, where the smallest standard deviation of the intensities is used for the estimation of noises [16]. All intensity-related parameters that we used in MZmine3 might change the peak list to some extent due to the different relative errors in intensity values.

#### 3.2.2 Comparison of Compound Lists among Intensity Groups with Different Relative Errors

The total number of detected compounds ranges from 3 (Met_10) to 9902(Met_04) (Table s4). After merging these repeated compounds, only about half of them were kept, called “types”. Three files (Met_10, Met_15 and Met_16) with a number of detected compounds less than 50 in all error groups were excluded from data statistics. The remaining 13 files contained a small fraction of unique compounds in the groups of 2×10^−4^ (average of 0.05%), 2×10^−3^ (average of 0.12%), 8×10^−3^ (average of 0.50%), 2×10^−2^ (average of 0.98%) (Fig. 4). However, the groups of 2×10^−1^ observed a decrease in the number of consensus compounds and had a roughly 7% of unique compounds. Thus, a file with an intensity relative error of less than 2×10^−2^ could result in a variance of less than 1% in compound identification compared to the precision-lossless files.

**Fig. 4.**
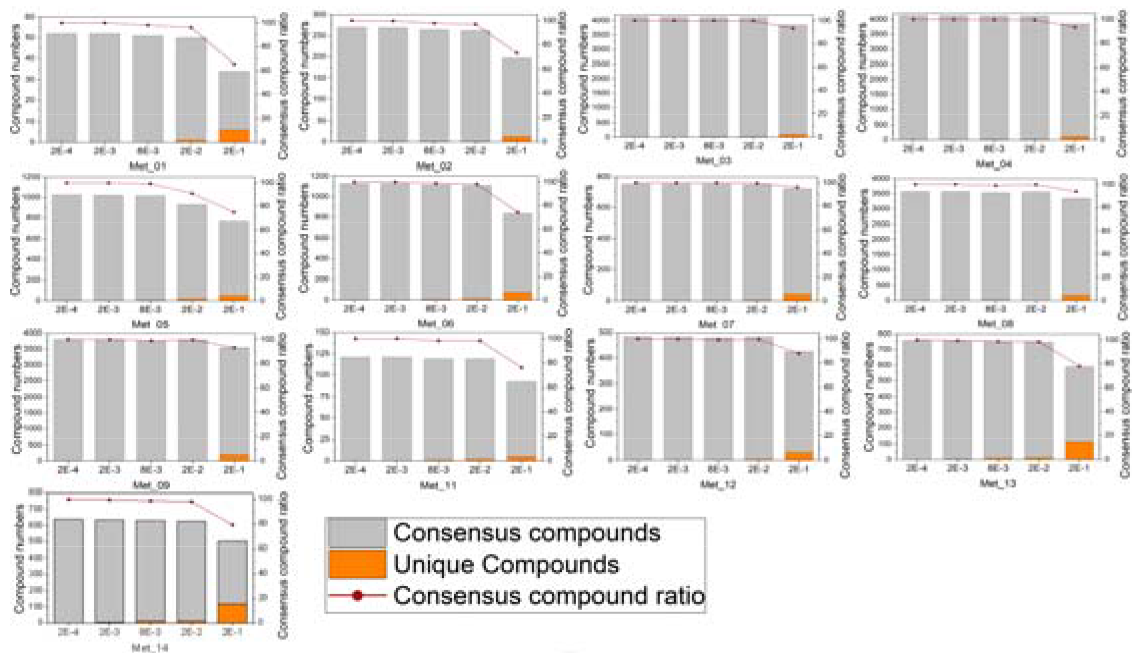
The number of total compounds and the percentage of consensus compounds in all five error groups across 13 mzML files with truncated intensity values.

### 3.3 Truncation Transform of m/z Values

Similar to intensities, we used the absolute error truncation transform for *m/z* values and set five groups (10^−5^, 2×10^−5^, 5×10^−5^, 10^−4^, 10^−3^).

#### 3.3.1 Comparison of Feature Lists among *m/z* Groups with Different Absolute Errors

The changes in the total numbers of detected features between blank and *m/z* error groups did not become significant until the error group (16 files) of 5×10^−5^ (p < 0.05) (Fig. s4 and Table s5). Next, we examined the proportion of consensus features in all detected features of a blank file and excluded the Met_10 file as the number of its detected peaks was too small to be statistically available. We then used the filtration window of *m/z* error <= 5ppm and RT shift <= 0.025min, resulting that consensus features took up over 99% of detected features in the first four groups (10^−5^ for 100%, 2×10^−5^ for 99.99%, 5×10^−5^ for 99.81%, 10^−4^ for 99.43%), but the group of 10^−3^ (92.39%) continued to lie below the line of 95% (Fig. 5). We further examined the intensity FC distribution of consensus features in all groups of files (Fig. s5).

**Fig. 5.**
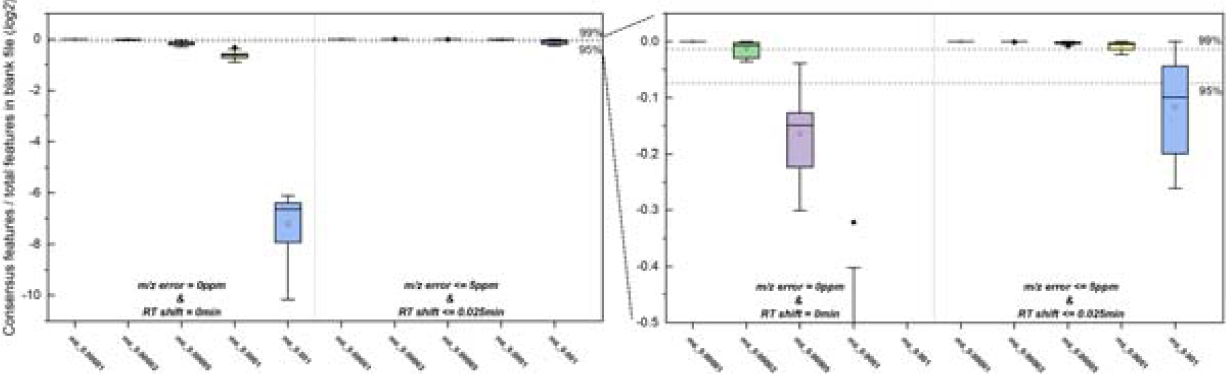
The ratio of consensus features to total features of blank files in each group with truncated *m/z* value under different filtration windows (*m/z* & RT error equals 0, *m/z* <= 5ppm & RT <= 0.025min). The ratio values were *log2* transformed. Met_10 was excluded due to the number of total features (n=10) was too small.

Overall, there are four groups (10^−5^, 2×10^−5^, 5×10^−5^, 10^−4^) with feature detection error under the threshold (1%). For significant features, a small fraction of less than 0.5% occurs in all groups, except about 0.65% (Met_14) in the group of 10^−2^.

To understand the variation generated from *m/z* error groups, tracking back to the data processing steps would be a preferred way. To discuss the effects of *m/z* values with errors on feature detection, we checked the ADAP peak-picking process again. For EIC building, all data points would be resorted from the highest intensity values to the lowest data and then be clustered to an EIC by the defined *m/z* accuracy (the parameter of scan-to-scan accuracy). In the study, *m/z* accuracy was set for TOF analyzers (0.005 *m/z* or 15 ppm) and orbitrap analyzers (0.005 *m/z* or 10 ppm) by the recommendation of MZmine3 documentation. The 15 ppm and 10 ppm at 500 *m/z* equal absolute errors of 0.0075 and 0.005 *m/z* (0.015 and 0.01 *m/z* for 15 and 10ppm at 1000 *m/z*), respectively. Therefore, the *m/z* with truncation transform at an absolute error of 10^−3^ is likely to be incorrectly included or excluded from the EIC-determining group. The errors from EICs would subsequently impact the peak detection step, where a certain part of misidentification features would not be filtered out as noise signals and eventually be kept as true features.

#### 3.3.2 Comparison of Compound Lists among *m/z* Groups with Different Absolute Errors

All the feature lists generated from files (except for Met_10, Met_15 and Met_16) with *m/z* errors have been processed by the online database search module of MZmine3. The distribution of consensus and unique compounds showed in Fig. 6. The number of unique compounds was increased by a noticeable degree from neglectable levels in the first four groups of 10^−5^ (0.03%), 2×10^−5^ (0.11%), 5×10^−5^ (0.52%), 10^−4^ (1.1%) to an average of 10.0% in the group of 10^−3^ (Table s5). Hence, the groups of 10^−5^, 2×10^−5^, 5×10^−5^ and 10^−4^ could control the UCR to about and below 1% of error threshold.

**Fig. 6.**
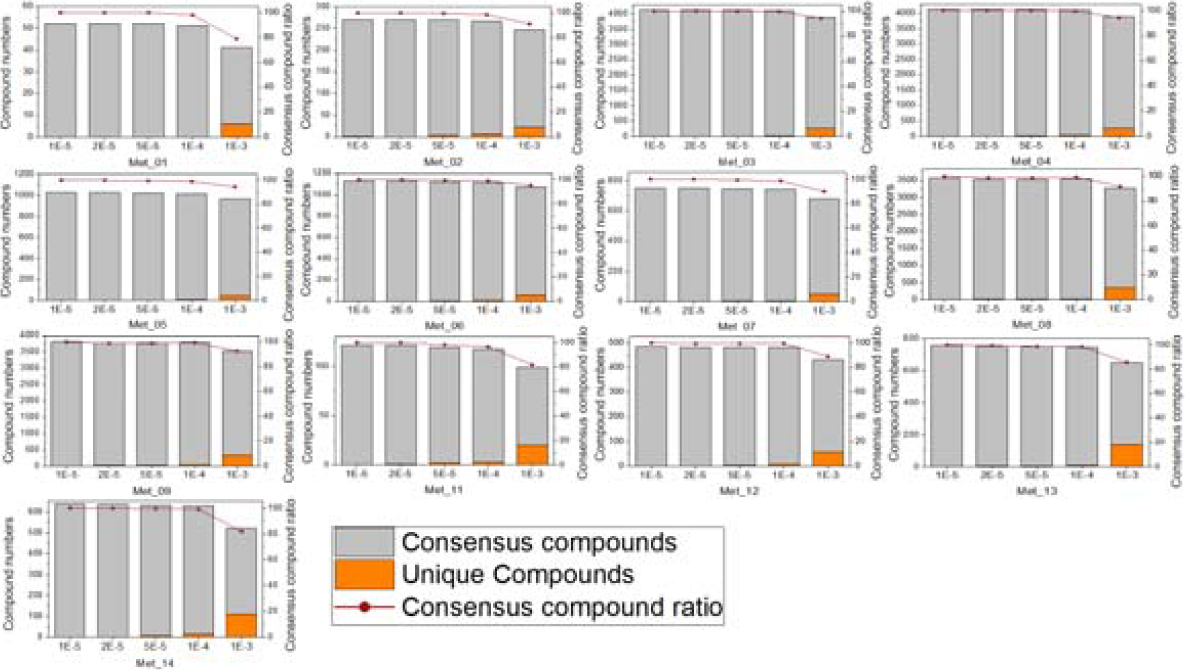
The number of total compounds and the percentage of cnsensus compounds in all five error groups across 13 mzML files with truncated *m/z* values

To conclude, we summarized the UFR, proportion of FC significant consensus features, and UCR in Table 12. For the intensity value, only two groups, 2×10^−4^ (0.04%) and 2×10^−3^(0.37%), had unique feature rates (UFR) of less than 1%, while the other groups had UFRs of varying degrees, such as 8×10^−3^ (1.55%), 2×10^−2^ (4.65%), and 2×10^−1^ (37.22%). We further checked the fraction of intensity FC significantly changed consensus features for each group, resulting in that all groups had a proportion of less than 1%. At the compound level, four of all groups (2×10^−4^ for 0.05%, 2×10^−3^ for 0.12%, 8×10^−3^ for 0.50%, 2×10^−2^ for 0.98%, 2×10^−1^ for 7%) with a variation of <1%. Therefore, we consider that 2×10^−2^ after annotation of the feature list (i.e., compounds detection) is considered as a preferred truncation error for lossy compression of intensity values. For *m/z* value, there are three groups (10^−5^ for 0%, 2×10^−5^ for 0.01%, 5×10^−5^ for 0.19%, 10^−4^ for 0.57%) showed UFRs below 1% in feature detection and the other one (10^−3^ for 7.61%) had higher UFRs. Furthermore, the proportion of significant FC consensus features was below 0.5% in all groups. In compound detection, four of all groups (10^−5^ for 0.03%, 2×10^−5^ for 0.11%, 5×10^−5^ for 0.52%, 10^−4^ for 1.10%, 10^−3^ for 10%) with a UCR below or around 1%. Thus, we reckon that 10^−4^ can be used as a suitable truncation error for the *m/z* values, since it is below and closet to 1% of error threshold in both feature and compound levels. Taken together, we proposed a maximum absolute error of 10^−4^ for *m/z* and a maximum relative error of 2×10^−2^ for intensity.

In this study, we only checked the untargeted metabolomics data in the MZmine3-based workflow, but the reproducibility of similar results on proteomics data and other software tools has not been examined yet. Moreover, the effects of precision loss on a downstream workflow of MZmine3, such as the feature based molecular network in GNPS [17] needed to be checked in our future step.

## 4. Conclusion

With the increase of MS data file size and volume, compressors specifically for MS data started to emerge. Among them, some lossy (near-lossless) compressors can reach a higher compression rate at the cost of precision losses for MS data (i.e., *m/z* and intensity). However, the extent to which the loss of precision at *m/z* and intensity have no negative impact on the subsequent data analysis has not been systematically discussed. In this study, sixteen vendor MS files were converted to mzML format via msConvert. We evaluated the difference among different combinations of storage precision in precision-lossless mzML files and the effects of precision losses on the data processing (feature detection and compound identification) steps via MZmine3.

First, we identified that mzML files with different combinations of storage precision at intensity and *m/z* data had minor variation on both feature and compound lists, with an average of 0.13%. We recognized that mzML files (precision-lossless) with both *m/z* and intensity encoded in 32-bit precision for the further evaluation of precision losses. We compared the results of feature and compound identification of precision-lossy files to precision-lossless files. A window of *m/z* error <= 5ppm and RT shift <= 0.025min was selected and used for consensus feature filtration. We eventually recommended a maximum absolute error of 10^−4^ for *m/z* and a maximum relative error of 2×10^−2^ for intensity, meeting the requirements of error threshold (1%). We further compared our recommendations to the recommended errors in existing near-losses compressors, such as MSNumpress (2×10^−9^ for *m/z* and 2×10^−4^ for intensity) and mspack (10^−4^ for *m/z* and 10^−2^ for intensity). According to our results, all error groups mentioned above would keep the variation of compound detection below 1%. However, we further extended the maximum relative error of intensity values to 2×10^−2^. We hope all results generated from our study could help researchers to make improvements to lossy compression algorithms in order to minimize the negative effects on downstream data processing, annotation and statistical steps.

## Supporting information

supplement tableS1 to figureS5 will be used for the link to the file on the preprint site

---

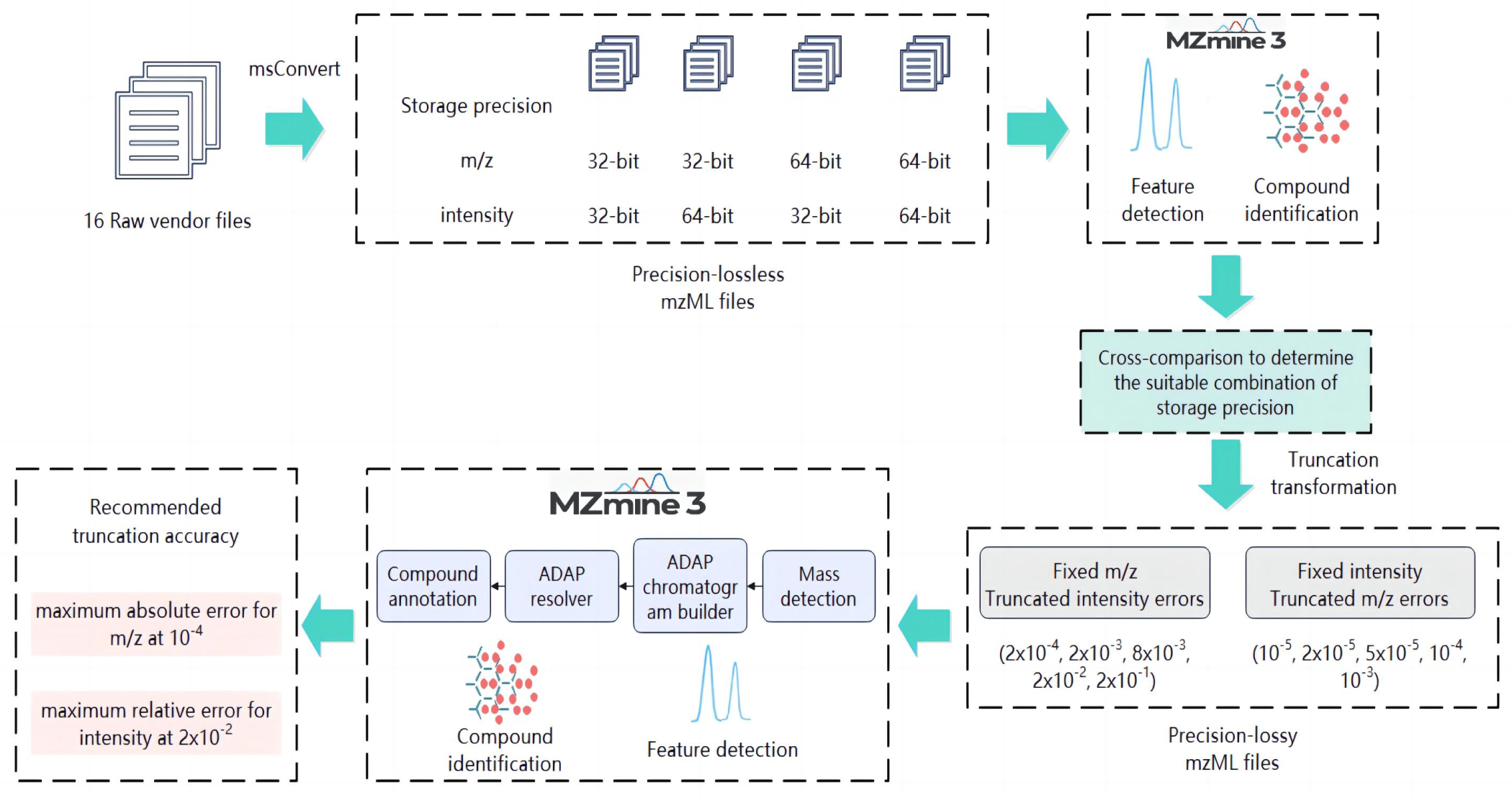

## References

1. Chen, Y., E.-M. Li, and L.-Y.J.M. Xu, Guide to Metabolomics Analysis: A Bioinformatics Workflow. 2022. 12(4): p. 357.

2. Schmid, R., et al., Integrative analysis of multimodal mass spectrometry data in MZmine 3. Nat Biotechnol, 2023.

3. Tsugawa, H., et al., MS-DIAL: data-independent MS/MS deconvolution for comprehensive metabolome analysis. Nat Methods, 2015. 12(6): p. 523–6.

4. Tautenhahn, R., et al., XCMS Online: a web-based platform to process untargeted metabolomic data. Anal Chem, 2012. 84(11): p. 5035–9.

5. Cui, L., H. Lu, and Y.H.J.M.s.r. Lee, Challenges and emergent solutions for LC-MS/MS based untargeted metabolomics in diseases. 2018. 37(6): p. 772–792.

6. Deutsch, E.W.J.M. and c. proteomics, File formats commonly used in mass spectrometry proteomics. 2012. 11(12): p. 1612–1621.

7. Martens, L., et al., mzML--a community standard for mass spectrometry data. Mol Cell Proteomics, 2011. 10(1): p. R110 000133.

8. Yang, R., X. Chen, and I. Ochoa, MassComp, a lossless compressor for mass spectrometry data. BMC Bioinformatics, 2019. 20(1): p. 368.

9. Teleman, J., et al., Numerical compression schemes for proteomics mass spectrometry data. Mol Cell Proteomics, 2014. 13(6): p. 1537–42.

10. Lu, M., et al., Aird: a computation-oriented mass spectrometry data format enables a higher compression ratio and less decoding time. BMC Bioinformatics, 2022. 23(1): p. 35.

11. Hanau, F., H. Rost, and I. Ochoa, mspack: efficient lossless and lossy mass spectrometry data compression. Bioinformatics, 2021.

12. Wang, J., et al., StackZDPD: a novel encoding scheme for mass spectrometry data optimized for speed and compression ratio. Sci Rep, 2022. 12(1): p. 5384.

13. Chambers, M.C., et al., A cross-platform toolkit for mass spectrometry and proteomics. Nat Biotechnol, 2012. 30(10): p. 918–20.

14. Tautenhahn, R., C. Bottcher, and S. Neumann, Highly sensitive feature detection for high resolution LC/MS. BMC Bioinformatics, 2008. 9: p. 504.

15. Myers, O.D., et al., Detailed Investigation and Comparison of the XCMS and MZmine 2 Chromatogram Construction and Chromatographic Peak Detection Methods for Preprocessing Mass Spectrometry Metabolomics Data. Anal Chem, 2017. 89(17): p. 8689–8695.

16. Myers, O.D., et al., One Step Forward for Reducing False Positive and False Negative Compound Identifications from Mass Spectrometry Metabolomics Data: New Algorithms for Constructing Extracted Ion Chromatograms and Detecting Chromatographic Peaks. Anal Chem, 2017. 89(17): p. 8696–8703.

17. Nothias, L.F., et al., Feature-based molecular networking in the GNPS analysis environment. Nat Methods, 2020. 17(9): p. 905–908.

