## supplement tableS1 to figureS5 will be used for the link to the file on the preprint site for "How much storage precision can be lost: Guidance for near-lossless compression of untargeted metabolomics mass spectrometry data"

**Table s1.** Raw MS vendor files used for evaluating the effects of storage precision on data processing steps. The information of file name, vendor instruments, data acquisition type and organisms in each file is included. Met_01 and 02 are from [1]. Met_03-06 are from [2]. Met_07 is from [3]. Met_08-09 are from [4]. Met_10 is from [5]. Met_11 is from [6]. Met_12 is from [7]. Met_13-14 are from [8]. Met_15-16 are from [9].

| File No. | Original File Name | Vendors Name | Instrument | Data Acquisition  Type | Organism (part) |
| --- | --- | --- | --- | --- | --- |
| Met_01 | Plasma3PlasmaDDA20-50.wiff | SCIEX | Triple TOF6600 | DDA | Homo sapiens (Blood Plasma) |
| Met_02 | PestMix-1-3-5-7Step1Plasma3SWATH20-50.wiff | SCIEX | Triple TOF6600 | DIA | Homo sapiens (Blood Plasma) |
| Met_03 | SA1.raw | Thermo | Q Exactive HF | DDA | Piper nigrum (Seed) |
| Met_04 | SB1.raw | Thermo | Q Exactive HF | DDA | Piper nigrum (Seed) |
| Met_05 | SampleA_1.wiff | SCIEX | Triple TOF6600 | DDA | Piper nigrum (Seed) |
| Met_06 | SampleB_1.wiff | SCIEX | Triple TOF6600 | DDA | Piper nigrum (Seed) |
| Met_07 | arunJ_20210318_o24433_pos_AIF_SOP4_empvec_1.raw | Thermo | Q Exactive | AIF | Saccharomyces cerevisiae (Whole Organism) |
| Met_08 | 04102020_Covid_TP_Mild_9030_1_P325_P_2.raw | Thermo | Q Exactive | DDA | Homo sapiens (Blood Plasma) |
| Met_09 | 23092020_Covid_TP_Severe_9030_2_P411_P_2.raw | Thermo | Q Exactive | DDA | Homo sapiens (Blood Plasma) |
| Met_10 | TFHN1.raw | Thermo | Q Exactive HF | DDA | Homo sapiens (Blood Plasma) |
| Met_11 | 180327_S1_LC02MS02_04_A12_ProlineBetaine.d | Agilent | 6550 Q-TOF | AIF | Homo sapiens (Urine) |
| Met_12 | 016_HEP_SOL1_LYS_neg_Spiking.raw | Thermo | Q Exactive HF | DDA | Homo sapiens (Hep-G2 cell) |
| Met_13 | CE04.raw | Thermo | Exactive Orbitrap | DDA | Homo sapiens (Urine) |
| Met_14 | CE05.raw | Thermo | Exactive Orbitrap | DDA | Homo sapiens (Urine) |
| Met_15 | Positive_000324.raw | Thermo | Q Exactive Plus | DDA | Saccharomyces cerevisiae (Whole Organism) |
| Met_16 | Negative_000333.raw | Thermo | Q Exactive Plus | DDA | Saccharomyces cerevisiae (Whole Organism) |

**Fig. s1. Comparison of detected features of different storage precision combinations across 16 files.** both *m/z* and intensity for 32-bit precision (Blue), both *m/z* and intensity for 64-bit precision (Green), *m/z* for 64-bit and intensity for 32-bit precision (Purple), *m/z* for 32-bit and intensity for 64-bit precision (Red).

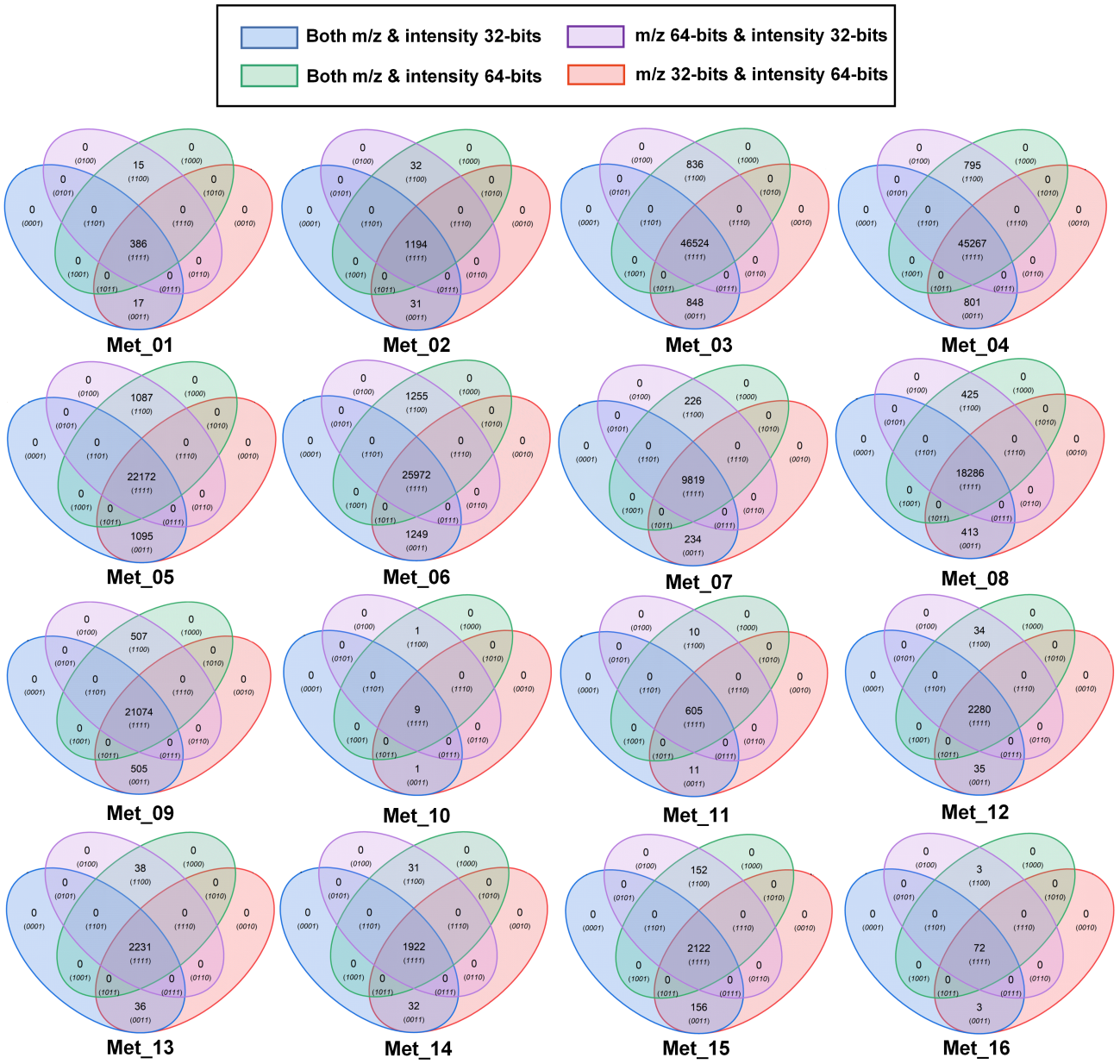

**Table s2.** The key parameters used for LC-MS data processing in MZmine3.

| **Module Name** | **Key Parameters** | **Contents** |
| --- | --- | --- |
| Mass detection | MS level | MS1 |
|  | Mass detector | Exact mass for profile data |
|  | Noise Level | 1.0E0 |
| ADAP chromatogram builder | Min group size of scans | 5 |
|  | Group intensity threshold | 5.0E2 |
|  | Min highest intensity | 5.0E4 |
|  | Scam to scan accuracy (*m/z*) | 0.0050 m/z or 15 ppm (TOF);  0.0050 m/z or 10ppm (Orbitrap) |
| ADAP resolver | S/N threshold | 10 |
|  | S/N estimator | Intensity window SN |
|  | min feature height | 10 |
|  | coefficient/area threshold | 110 |
|  | Peak duration range | 0.00-10.00 |
|  | RT wavelet range | 0.00-0.10 |
| Join aligner | *m/z* tolerance | 0.001 *m/z* or 5 ppm |
|  | Weight for *m/z* | 5 |
|  | Retention time tolerance | 0.01 |
|  | Weight for RT | 5 |

**Table s3.** The summary of compound identification results in all precision-lossless mzML files.

|  |  | Online Compound Database Search | | | | | | |
| --- | --- | --- | --- | --- | --- | --- | --- | --- |
|  |  | Used Database | Detected Total Compounds | Types of Compounds | Consensus Compounds | Unique Compounds | Unique Compound  Rate % | Compounds Annotated by Unique Features |
| Met_01 | mz32 | HMDB | 124 | 52 | 51 | 1 | 1.92% | 1(1*) |
|  | mz64 |  | 123 | 51 |  | NA | NA | 0 |
| Met_02 | mz32 | HMDB | 624 | 270 | 270 | NA | NA | 0 |
|  | mz64 |  | 625 | 271 |  | 1 | 0.37% | 0 |
| Met_03 | mz32 | KEGG | 9765 | 4124 | 4123 | 1 | 0.02% | 17 |
|  | mz64 |  | 9766 | 4125 |  | 2 | 0.05% | 17(1*) |
| Met_04 | mz32 | KEGG | 9902 | 4134 | 4128 | 6 | 0.15% | 24(4*) |
|  | mz64 |  | 9901 | 4132 |  | 4 | 0.10% | 23(2*) |
| Met_05 | mz32 | KEGG | 7042 | 1024 | 1023 | 1 | 0.10% | 16 |
|  | mz64 |  | 7049 | 1024 |  | 1 | 0.10% | 21 |
| Met_06 | mz32 | KEGG | 8513 | 1128 | 1125 | 3 | 0.27% | 23 |
|  | mz64 |  | 8522 | 1130 |  | 5 | 0.44% | 18(2*) |
| Met_07 | mz32 | KEGG | 1239 | 752 | 751 | 1 | 0.13% | 0 |
|  | mz64 |  | 1238 | 751 |  | NA | NA | 0 |
| Met_08 | mz32 | HMDB | 6367 | 3568 | 3564 | 4 | 0.11% | 5(3*) |
|  | mz64 |  | 6375 | 3568 |  | 4 | 0.11% | 13(3*) |
| Met_09 | mz32 | HMDB | 7392 | 3815 | 3812 | 3 | 0.08% | 10(2*) |
|  | mz64 |  | 7398 | 3816 |  | 4 | 0.10% | 16(2*) |
| Met_10 | mz32 | HMDB | 3 | 3 | 3 | NA | NA | 0 |
|  | mz64 |  | 3 | 3 |  | NA | NA | 0 |
| Met_11 | mz32 | HMDB | 229 | 121 | 121 | NA | NA | 1 |
|  | mz64 |  | 229 | 122 |  | 1 | 0.82% | 0 |
| Met_12 | mz32 | HMDB | 621 | 485 | 485 | NA | NA | 1 |
|  | mz64 |  | 620 | 485 |  | NA | NA | 0 |
| Met_13 | mz32 | HMDB | 1096 | 759 | 759 | NA | NA | 0 |
|  | mz64 |  | 1097 | 759 |  | NA | NA | 0 |
| Met_14 | mz32 | HMDB | 922 | 638 | 638 | NA | NA | 0 |
|  | mz64 |  | 923 | 639 |  | 1 | 0.16% | 0 |
| Met_15 | mz32 | YMDB | 26 | 20 | 19 | 1 | 5.00% | 1(1*) |
|  | mz64 |  | 25 | 19 |  | NA | NA | 0 |
| Met_16 | mz32 | KEGG | 9 | 8 | 8 | NA | NA | 0 |
|  | mz64 |  | 9 | 8 |  | NA | NA | 0 |

* means the number of unique compounds annotated by unique features

NA means zero

**Fig. s2.** The number of consensus features detected in all groups (Blank, 2x10^-4^, 2x10^-3^, 8x10^-3^, 2x10^-2^, 2x10^-1^) across all files (Met_01 – Met_16) with truncated intensities.

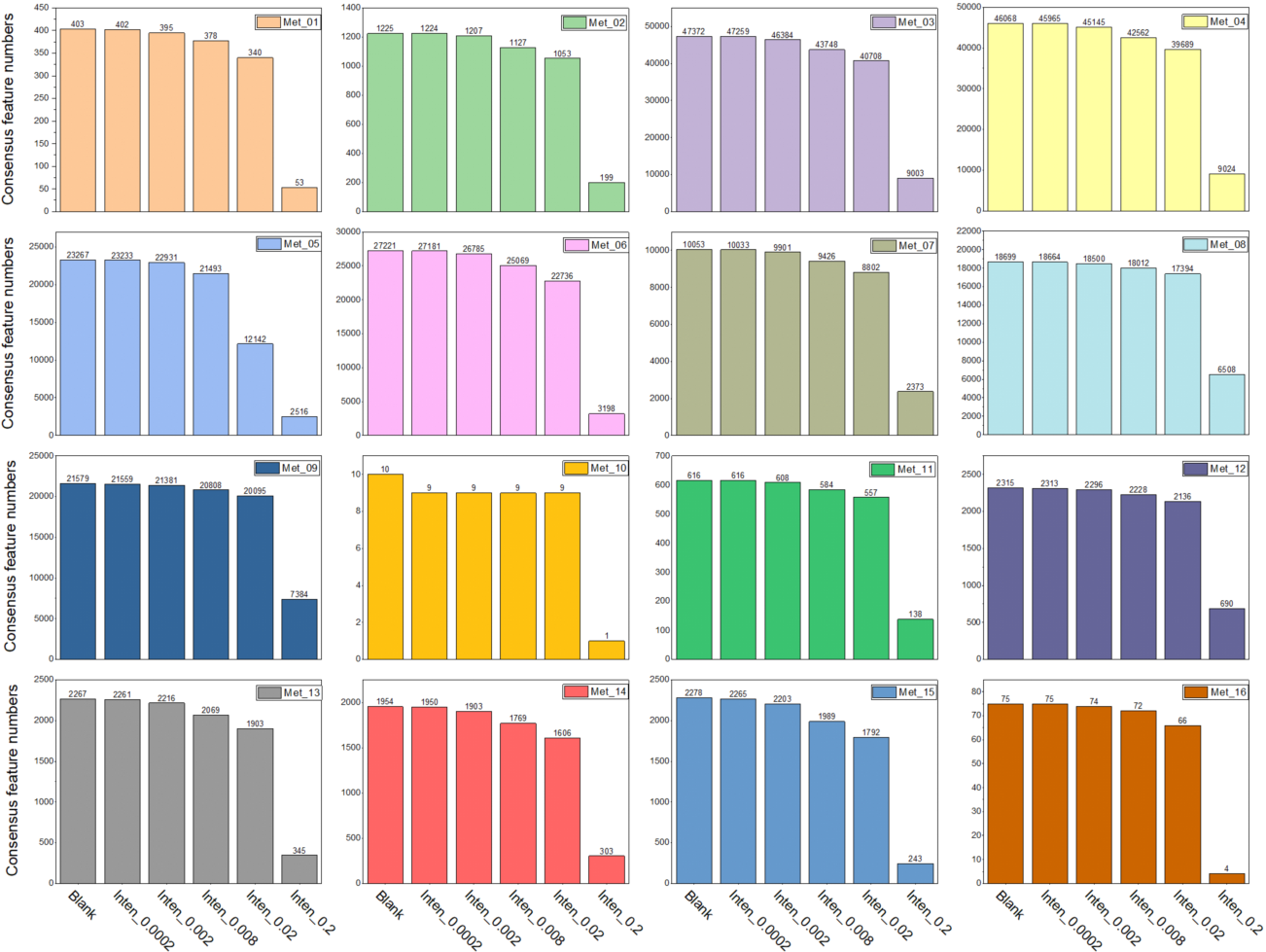

**Fig. s3. The summary of intensities fold-change (FC) of consensus features in all groups of files with truncated intensities.** The box chart shows *Log_2_* transformed FC values in all groups (2x10^-4^, 2x10^-3^, 8x10^-3^, 2x10^-2^, 2x10^-1^).

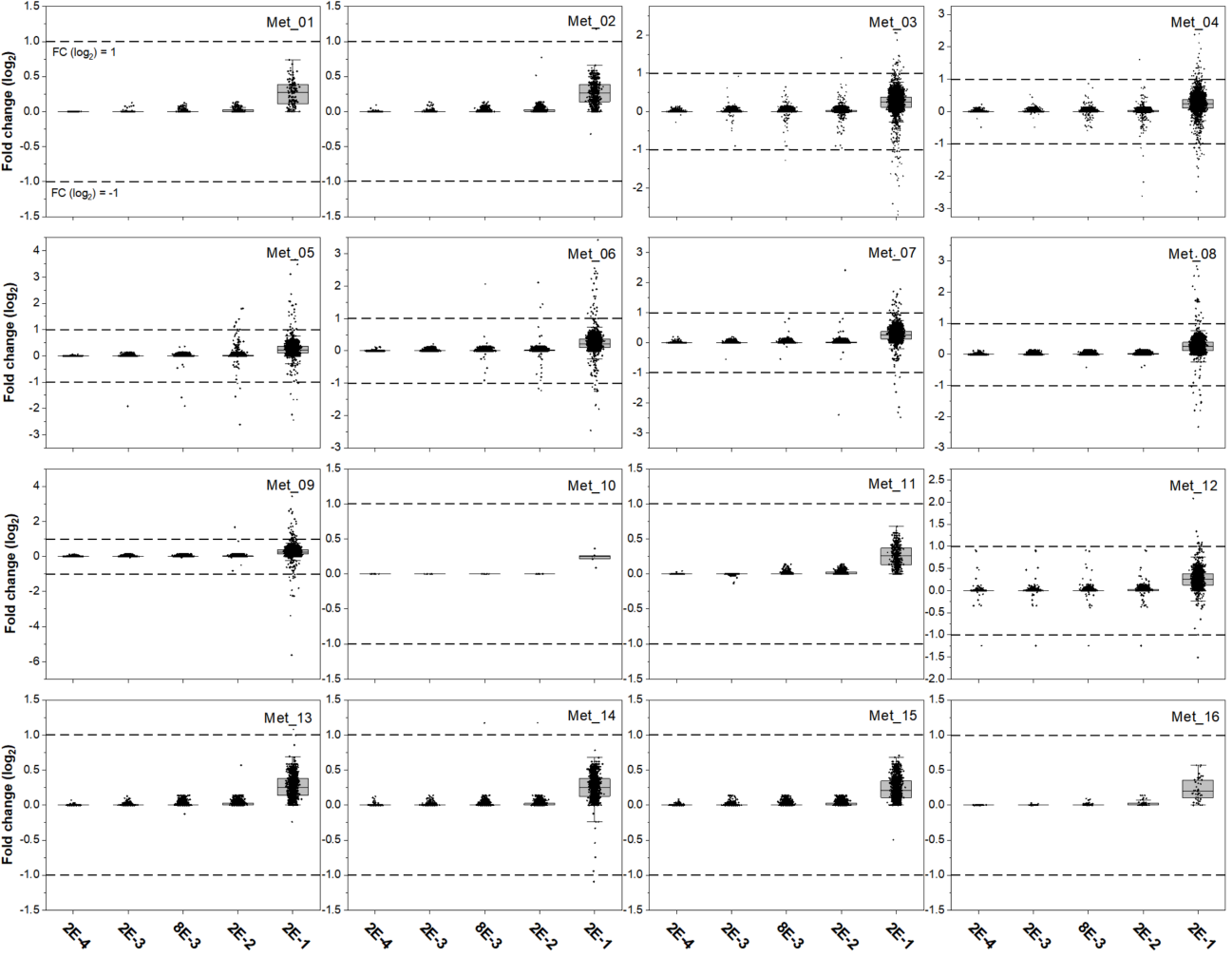

Consensus features shared the m/z and RT value, while the difference between their intensities was unknown and could impact the compound identification in the downstream of data analysis. For the filtered consensus features, we calculated the fold-change (FC) of their intensities between blank files and error groups (Figure6). We defined significant features with FC >2 (upper dash line in Figure6) or <0.5 (lower dash line in Figure6) and less significant features (0.5< FC <2, except FC=1). All groups have a small fraction of significant features (less than 0.5%), except about 0.63% (Met_12) and 0.55% (Met_06) in the group of 2x10-1. The proportion of less significant features in consensus features for groups were vary (2x10-4 for <1%, 2x10-3 for <5%, 8x10-3 for about 20%, 2x10-2 for about 40%, 2x10-1 for about 95%). Therefore, the fraction of significant features was considered in a controllable level in all groups (mostly less than 0.5%). However, the fraction of less significant features suggested that 2x10-4 and 2x10-3 seems to be preferred error groups with a number below 5%.

**Table s4. The summary of detected features and compounds in mzML files with truncated intensity values**. Consensus features (unfiltered) are features in error groups overlapping with blank files. Unique features are features only detected in error groups. Types means the compounds after removing repeated ones.

|  |  | Detected Features | | | Detected compounds | | | | |
| --- | --- | --- | --- | --- | --- | --- | --- | --- | --- |
|  |  | Total | Consensus features | Unique  features | Total | Types | Consensus compounds | Unique compounds | Annotated by Unique Features |
| Met_01 | Blank | 403 |  |  | 124 | 52 |  |  |  |
|  | Inten_2E-4 | 403 | 402 | 1 | 124 | 52 | 52 | 0 | 0 |
|  | Inten_2E-3 | 403 | 395 | 8 | 124 | 52 | 52 | 0 | 0 |
|  | Inten_8E-3 | 402 | 378 | 24 | 121 | 51 | 51 | 0 | 0 |
|  | Inten_2E-2 | 403 | 340 | 63 | 123 | 51 | 50 | 1 | 5(1*) |
|  | Inten_2E-1 | 333 | 53 | 280 | 79 | 40 | 34 | 6 | 22（5*） |
| Met_02 | Blank | 1225 |  |  | 624 | 270 |  |  |  |
|  | Inten_2E-4 | 1225 | 1224 | 1 | 625 | 271 | 270 | 1 | 0 |
|  | Inten_2E-3 | 1223 | 1207 | 16 | 623 | 270 | 269 | 1 | 0 |
|  | Inten_8E-3 | 1209 | 1127 | 82 | 617 | 265 | 264 | 1 | 12 |
|  | Inten_2E-2 | 1209 | 1053 | 156 | 616 | 263 | 262 | 1 | 23 |
|  | Inten_2E-1 | 995 | 199 | 796 | 503 | 210 | 197 | 13 | 118(11*) |
| Met_03 | Blank | 47372 |  |  | 9765 | 4124 |  |  |  |
|  | Inten_2E-4 | 47370 | 47259 | 111 | 9766 | 4124 | 4124 | 0 | 3 |
|  | Inten_2E-3 | 47381 | 46384 | 997 | 9767 | 4122 | 4122 | 0 | 2 |
|  | Inten_8E-3 | 47352 | 43748 | 3604 | 9765 | 4125 | 4117 | 8 | 64(5*) |
|  | Inten_2E-2 | 47349 | 40708 | 6641 | 9776 | 4120 | 4108 | 12 | 122(7*) |
|  | Inten_2E-1 | 47727 | 9003 | 38724 | 10007 | 3942 | 3847 | 95 | 964(86*) |
| Met_04 | Blank | 46068 |  |  | 9902 | 4134 |  |  |  |
|  | Inten_2E-4 | 46064 | 45965 | 99 | 9902 | 4135 | 4134 | 1 | 2 |
|  | Inten_2E-3 | 46065 | 45145 | 920 | 9903 | 4136 | 4131 | 5 | 16(3*) |
|  | Inten_8E-3 | 46062 | 42562 | 3500 | 9900 | 4133 | 4120 | 13 | 65(10*) |
|  | Inten_2E-2 | 46081 | 39689 | 6392 | 9900 | 4132 | 4115 | 17 | 119(12*) |
|  | Inten_2E-1 | 46371 | 9024 | 37347 | 10069 | 3976 | 3858 | 118 | 1002(97*) |
| Met_05 | Blank | 23267 |  |  | 7042 | 1024 |  |  |  |
|  | Inten_2E-4 | 23273 | 23233 | 40 | 7043 | 1024 | 1024 | 0 | 2 |
|  | Inten_2E-3 | 23250 | 22931 | 319 | 7031 | 1022 | 1022 | 0 | 19 |
|  | Inten_8E-3 | 23230 | 21493 | 1737 | 7050 | 1020 | 1016 | 4 | 133(4*) |
|  | Inten_2E-2 | 21353 | 12142 | 9211 | 6639 | 948 | 930 | 18 | 497(18*) |
|  | Inten_2E-1 | 18855 | 2516 | 16339 | 5650 | 817 | 766 | 51 | 707(49*) |
| Met_06 | Blank | 27221 |  |  | 8513 | 1128 |  |  |  |
|  | Inten_2E-4 | 27231 | 27181 | 50 | 8520 | 1129 | 1128 | 1 | 0 |
|  | Inten_2E-3 | 27176 | 26785 | 391 | 8521 | 1130 | 1128 | 2 | 20(1*) |
|  | Inten_8E-3 | 27079 | 25069 | 2010 | 8484 | 1124 | 1116 | 8 | 170(8*) |
|  | Inten_2E-2 | 27002 | 22736 | 4266 | 8408 | 1123 | 1109 | 14 | 294(14*) |
|  | Inten_2E-1 | 22641 | 3198 | 19443 | 6827 | 910 | 836 | 75 | 831(75*) |
| Met_07 | Blank | 10053 |  |  | 1239 | 752 |  |  |  |
|  | Inten_2E-4 | 10054 | 10033 | 21 | 1239 | 752 | 752 | 0 | 0 |
|  | Inten_2E-3 | 10058 | 9901 | 157 | 1240 | 751 | 751 | 0 | 1 |
|  | Inten_8E-3 | 10051 | 9426 | 625 | 1241 | 751 | 751 | 0 | 6 |
|  | Inten_2E-2 | 10062 | 8802 | 1260 | 1243 | 752 | 748 | 4 | 21(3*) |
|  | Inten_2E-1 | 10756 | 2373 | 8383 | 1484 | 766 | 722 | 44 | 167(35*) |
| Met_08 | Blank | 18699 |  |  | 6367 | 3568 |  |  |  |
|  | Inten_2E-4 | 18704 | 18664 | 40 | 6368 | 3569 | 3568 | 1 | 2(1*) |
|  | Inten_2E-3 | 18712 | 18500 | 212 | 6374 | 3572 | 3568 | 4 | 9(4*) |
|  | Inten_8E-3 | 18701 | 18012 | 689 | 6318 | 3543 | 3533 | 10 | 20(5*) |
|  | Inten_2E-2 | 18718 | 17394 | 1324 | 6369 | 3565 | 3560 | 5 | 35(5*) |
|  | Inten_2E-1 | 20240 | 6508 | 13732 | 6925 | 3520 | 3345 | 175 | 581(148*) |
| Met_09 | Blank | 21579 |  |  | 7392 | 3815 |  |  |  |
|  | Inten_2E-4 | 21580 | 21559 | 21 | 7392 | 3815 | 3815 | 0 | 1 |
|  | Inten_2E-3 | 21580 | 21381 | 199 | 7396 | 3817 | 3815 | 2 | 10(2*) |
|  | Inten_8E-3 | 21558 | 20808 | 750 | 7321 | 3781 | 3774 | 7 | 23(4*) |
|  | Inten_2E-2 | 21558 | 20095 | 1463 | 7402 | 3815 | 3806 | 9 | 44(9*) |
|  | Inten_2E-1 | 22829 | 7384 | 15445 | 7910 | 3750 | 3553 | 197 | 697(174*) |
| Met_10 | Blank | 10 |  |  | 3 | 3 |  |  |  |
|  | Inten_2E-4 | 9 | 9 | 0 | 3 | 3 | 3 | 0 | 0 |
|  | Inten_2E-3 | 9 | 9 | 0 | 3 | 3 | 3 | 0 | 0 |
|  | Inten_8E-3 | 9 | 9 | 0 | 3 | 3 | 3 | 0 | 0 |
|  | Inten_2E-2 | 9 | 9 | 0 | 3 | 3 | 3 | 0 | 0 |
|  | Inten_2E-1 | 8 | 1 | 7 | 4 | 4 | 3 | 1 | 2 |
| Met_11 | Blank | 616 |  |  | 229 | 121 |  |  |  |
|  | Inten_2E-4 | 616 | 616 | 0 | 229 | 121 | 121 | 0 | 0 |
|  | Inten_2E-3 | 616 | 608 | 8 | 229 | 121 | 121 | 0 | 0 |
|  | Inten_8E-3 | 608 | 584 | 24 | 224 | 120 | 119 | 1 | 1 |
|  | Inten_2E-2 | 613 | 557 | 56 | 228 | 121 | 119 | 2 | 5(1*) |
|  | Inten_2E-1 | 612 | 138 | 474 | 218 | 97 | 92 | 5 | 29(5*) |
| Met_12 | Blank | 2315 |  |  | 621 | 485 |  |  |  |
|  | Inten_2E-4 | 2315 | 2313 | 2 | 621 | 485 | 485 | 0 | 0 |
|  | Inten_2E-3 | 2315 | 2296 | 19 | 620 | 484 | 484 | 0 | 0 |
|  | Inten_8E-3 | 2317 | 2228 | 89 | 616 | 482 | 481 | 1 | 1(1*) |
|  | Inten_2E-2 | 2321 | 2136 | 185 | 622 | 486 | 483 | 3 | 8(3*) |
|  | Inten_2E-1 | 2142 | 690 | 1452 | 572 | 458 | 426 | 32 | 67(27*) |
| Met_13 | Blank | 2267 |  |  | 1096 | 759 |  |  |  |
|  | Inten_2E-4 | 2266 | 2261 | 5 | 1096 | 759 | 759 | 0 | 0 |
|  | Inten_2E-3 | 2263 | 2216 | 47 | 1094 | 758 | 755 | 3 | 9(2*) |
|  | Inten_8E-3 | 2272 | 2069 | 203 | 1099 | 760 | 749 | 11 | 42(9*) |
|  | Inten_2E-2 | 2277 | 1903 | 374 | 1105 | 761 | 748 | 13 | 69(12*) |
|  | Inten_2E-1 | 2146 | 345 | 1801 | 1061 | 704 | 590 | 114 | 285(103*) |
| Met_14 | Blank | 1954 |  |  | 922 | 638 |  |  |  |
|  | Inten_2E-4 | 1952 | 1950 | 2 | 922 | 639 | 638 | 1 | 1(1*) |
|  | Inten_2E-3 | 1954 | 1903 | 51 | 922 | 638 | 636 | 2 | 7(2*) |
|  | Inten_8E-3 | 1961 | 1769 | 192 | 927 | 641 | 631 | 10 | 19(8*) |
|  | Inten_2E-2 | 1962 | 1606 | 356 | 926 | 638 | 626 | 12 | 44(11*) |
|  | Inten_2E-1 | 1889 | 303 | 1586 | 946 | 621 | 505 | 116 | 261(109*) |
| Met_15 | Blank | 2278 |  |  | 26 | 20 |  |  |  |
|  | Inten_2E-4 | 2278 | 2265 | 13 | 26 | 20 | 20 | 0 | 0 |
|  | Inten_2E-3 | 2282 | 2203 | 79 | 26 | 20 | 20 | 0 | 0 |
|  | Inten_8E-3 | 2293 | 1989 | 304 | 26 | 20 | 20 | 0 | 0 |
|  | Inten_2E-2 | 2305 | 1792 | 513 | 27 | 22 | 20 | 2 | 2(2*) |
|  | Inten_2E-1 | 2743 | 243 | 2500 | 28 | 19 | 15 | 4 | 11(4*) |
| Met_16 | Blank | 75 |  |  | 9 | 8 |  |  |  |
|  | Inten_2E-4 | 75 | 75 | 0 | 9 | 8 | 8 | 0 | 0 |
|  | Inten_2E-3 | 75 | 74 | 1 | 9 | 8 | 8 | 0 | 0 |
|  | Inten_8E-3 | 76 | 72 | 4 | 9 | 8 | 8 | 0 | 0 |
|  | Inten_2E-2 | 78 | 66 | 12 | 9 | 8 | 8 | 0 | 0 |
|  | Inten_2E-1 | 122 | 4 | 118 | 20 | 19 | 8 | 11 | 12(11*) |

* means the number of unique compounds annotated by unique features

**Fig. s4.** The number of consensus features detected in all groups (Blank, 10^-5^, 2x10^-5^, 5x10^-5^, 10^-4^, 10^-3^) across all files (Met_01 – Met_16) with truncated *m/z* data.

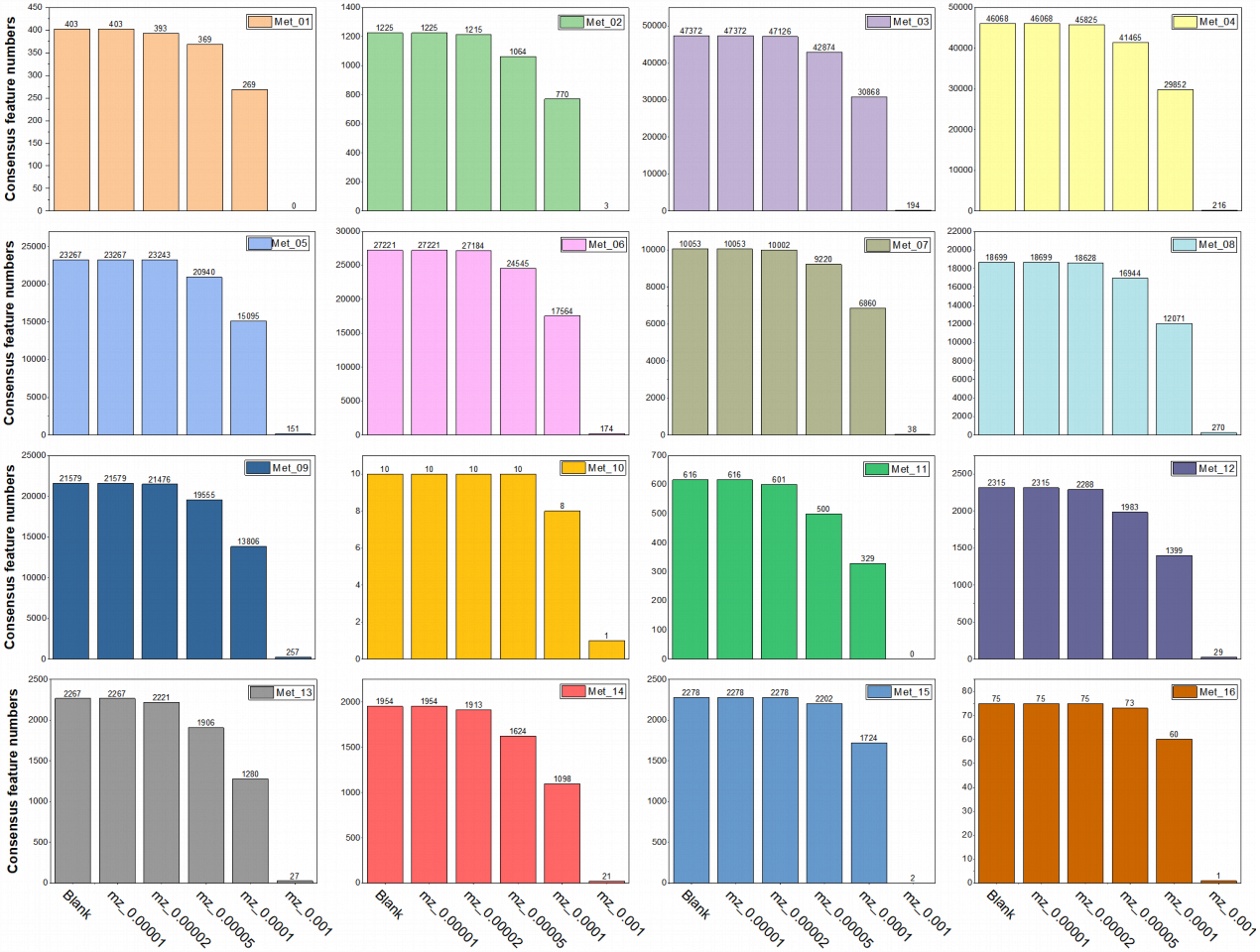

**Table s5. The summary of detected features and compounds in mzML files with truncated *m/z* values.** Consensus features (unfiltered) are features in error groups overlapping with blank files. Unique features are features only detected in error groups. Types means the compounds after removing repeated ones.

|  |  | Detected Features | | | Detected compounds | | | | |
| --- | --- | --- | --- | --- | --- | --- | --- | --- | --- |
|  |  | Total | True positive | False positive | Total | Types | True positive | False positive | Annotated by False Features |
| Met_01 | Blank | 403 |  |  | 124 | 52 |  |  |  |
|  | mz_1E-5 | 403 | 403 | 0 | 124 | 52 | 52 | 0 | 0 |
|  | mz_2E-5 | 403 | 393 | 10 | 124 | 52 | 52 | 0 | 2 |
|  | mz_5E-5 | 403 | 369 | 34 | 124 | 52 | 52 | 0 | 2 |
|  | mz_1E-4 | 402 | 269 | 133 | 123 | 51 | 51 | 0 | 4 |
|  | mz_1E-3 | 395 | 0 | 395 | 116 | 47 | 41 | 6 | 40(5*) |
| Met_02 | Blank | 1225 |  |  | 624 | 270 |  |  |  |
|  | mz_1E-5 | 1225 | 1225 | 0 | 625 | 271 | 270 | 1 | 0 |
|  | mz_2E-5 | 1225 | 1215 | 10 | 624 | 270 | 270 | 0 | 0 |
|  | mz_5E-5 | 1227 | 1064 | 163 | 626 | 272 | 269 | 3 | 3 |
|  | mz_1E-4 | 1223 | 770 | 453 | 624 | 272 | 266 | 6 | 2 |
|  | mz_1E-3 | 1211 | 3 | 1208 | 619 | 269 | 246 | 23 | 248(22*) |
| Met_03 | Blank | 47372 |  |  | 9765 | 4124 |  |  |  |
|  | mz_1E-5 | 47372 | 47372 | 0 | 9765 | 4124 | 4124 | 0 | 0 |
|  | mz_2E-5 | 47376 | 47126 | 250 | 9765 | 4124 | 4124 | 0 | 0 |
|  | mz_5E-5 | 47362 | 42874 | 4488 | 9763 | 4123 | 4118 | 5 | 18(2*) |
|  | mz_1E-4 | 47359 | 30868 | 16491 | 9767 | 4128 | 4107 | 21 | 40(8*) |
|  | mz_1E-3 | 48257 | 194 | 48063 | 9916 | 4148 | 3885 | 263 | 3300(233*) |
| Met_04 | Blank | 46068 |  |  | 9902 | 4134 |  |  |  |
|  | mz_1E-5 | 46068 | 46068 | 0 | 9902 | 4134 | 4134 | 0 | 0 |
|  | mz_2E-5 | 46071 | 45825 | 246 | 9903 | 4134 | 4134 | 0 | 0 |
|  | mz_5E-5 | 46066 | 41465 | 4601 | 9898 | 4137 | 4128 | 9 | 16(1*) |
|  | mz_1E-4 | 46092 | 29852 | 16240 | 9877 | 4141 | 4116 | 25 | 38(6*) |
|  | mz_1E-3 | 46908 | 216 | 46692 | 9968 | 4157 | 3895 | 262 | 3303(238*) |
| Met_05 | Blank | 23267 |  |  | 7042 | 1024 |  |  |  |
|  | mz_1E-5 | 23267 | 23267 | 0 | 7042 | 1024 | 1024 | 0 | 0 |
|  | mz_2E-5 | 23267 | 23243 | 24 | 7042 | 1024 | 1024 | 0 | 0 |
|  | mz_5E-5 | 23228 | 20940 | 2288 | 7044 | 1021 | 1020 | 1 | 45(1*) |
|  | mz_1E-4 | 23227 | 15095 | 8132 | 7044 | 1020 | 1014 | 6 | 119(5*) |
|  | mz_1E-3 | 23396 | 151 | 23245 | 7157 | 1010 | 969 | 41 | 946(35*) |
| Met_06 | Blank | 27221 |  |  | 8513 | 1128 |  |  |  |
|  | mz_1E-5 | 27221 | 27221 | 0 | 8513 | 1128 | 1128 | 0 | 0 |
|  | mz_2E-5 | 27220 | 27184 | 36 | 8512 | 1128 | 1128 | 0 | 0 |
|  | mz_5E-5 | 27209 | 24545 | 2664 | 8497 | 1128 | 1122 | 6 | 55(3*) |
|  | mz_1E-4 | 27231 | 17564 | 9667 | 8507 | 1127 | 1117 | 10 | 133(6*) |
|  | mz_1E-3 | 27166 | 174 | 26992 | 8678 | 1129 | 1075 | 54 | 1071(51*) |
| Met_07 | Blank | 10053 |  |  | 1239 | 752 |  |  |  |
|  | mz_1E-5 | 10053 | 10053 | 0 | 1239 | 752 | 752 | 0 | 0 |
|  | mz_2E-5 | 10053 | 10002 | 51 | 1240 | 752 | 752 | 0 | 0 |
|  | mz_5E-5 | 10053 | 9220 | 833 | 1236 | 750 | 747 | 3 | 0 |
|  | mz_1E-4 | 10056 | 6860 | 3196 | 1233 | 747 | 742 | 5 | 1(1*) |
|  | mz_1E-3 | 10098 | 38 | 10060 | 1213 | 730 | 678 | 52 | 559(48*) |
| Met_08 | Blank | 18699 |  |  | 6367 | 3568 |  |  |  |
|  | mz_1E-5 | 18699 | 18699 | 0 | 6367 | 3568 | 3568 | 0 | 0 |
|  | mz_2E-5 | 18700 | 18628 | 72 | 6315 | 3542 | 3534 | 8 | 1(1*) |
|  | mz_5E-5 | 18705 | 16944 | 1761 | 6315 | 3543 | 3528 | 15 | 10(2*) |
|  | mz_1E-4 | 18696 | 12071 | 6625 | 6361 | 3567 | 3545 | 22 | 22(4*) |
|  | mz_1E-3 | 19412 | 270 | 19142 | 6642 | 3635 | 3279 | 356 | 2600(318*) |
| Met_09 | Blank | 21579 |  |  | 7392 | 3815 |  |  |  |
|  | mz_1E-5 | 21579 | 21579 | 0 | 7392 | 3815 | 3815 | 0 | 0 |
|  | mz_2E-5 | 21578 | 21476 | 102 | 7327 | 3783 | 3779 | 4 | 0 |
|  | mz_5E-5 | 21587 | 19555 | 2032 | 7329 | 3786 | 3772 | 14 | 6(2*) |
|  | mz_1E-4 | 21585 | 13806 | 7779 | 7387 | 3820 | 3789 | 31 | 26(7*) |
|  | mz_1E-3 | 22213 | 257 | 21956 | 7538 | 3859 | 3531 | 328 | 2704(283*) |
| Met_10 | Blank | 10 |  |  | 3 | 3 |  |  |  |
|  | mz_1E-5 | 10 | 10 | 0 | 3 | 3 | 3 | 0 | 0 |
|  | mz_2E-5 | 10 | 10 | 0 | 3 | 3 | 3 | 0 | 0 |
|  | mz_5E-5 | 10 | 10 | 0 | 3 | 3 | 3 | 0 | 0 |
|  | mz_1E-4 | 10 | 8 | 2 | 3 | 3 | 3 | 0 | 0 |
|  | mz_1E-3 | 9 | 1 | 8 | 3 | 3 | 3 | 0 | 0 |
| Met_11 | Blank | 616 |  |  | 229 | 121 |  |  |  |
|  | mz_1E-5 | 616 | 616 | 0 | 229 | 121 | 121 | 0 | 0 |
|  | mz_2E-5 | 616 | 601 | 15 | 231 | 122 | 121 | 1 | 0 |
|  | mz_5E-5 | 616 | 500 | 116 | 229 | 121 | 119 | 2 | 2 |
|  | mz_1E-4 | 615 | 329 | 286 | 229 | 120 | 117 | 3 | 5 |
|  | mz_1E-3 | 613 | 0 | 613 | 213 | 119 | 99 | 20 | 108(18*) |
| Met_12 | Blank | 2315 |  |  | 621 | 485 |  |  |  |
|  | mz_1E-5 | 2315 | 2315 | 0 | 621 | 485 | 485 | 0 | 0 |
|  | mz_2E-5 | 2315 | 2288 | 27 | 618 | 482 | 482 | 0 | 0 |
|  | mz_5E-5 | 2316 | 1983 | 333 | 620 | 484 | 482 | 2 | 2 |
|  | mz_1E-4 | 2314 | 1399 | 915 | 630 | 491 | 483 | 8 | 5(2*) |
|  | mz_1E-3 | 2338 | 29 | 2309 | 627 | 489 | 432 | 57 | 398(52*) |
| Met_13 | Blank | 2267 |  |  | 1096 | 759 |  |  |  |
|  | mz_1E-5 | 2267 | 2267 | 0 | 1096 | 759 | 759 | 0 | 0 |
|  | mz_2E-5 | 2269 | 2221 | 48 | 1095 | 756 | 755 | 1 | 2 |
|  | mz_5E-5 | 2273 | 1906 | 367 | 1089 | 750 | 749 | 1 | 8 |
|  | mz_1E-4 | 2251 | 1280 | 971 | 1085 | 752 | 747 | 5 | 19 |
|  | mz_1E-3 | 2444 | 27 | 2417 | 1137 | 786 | 649 | 137 | 735(135*) |
| Met_14 | Blank | 1954 |  |  | 922 | 638 |  |  |  |
|  | mz_1E-5 | 1954 | 1954 | 0 | 922 | 638 | 638 | 0 | 0 |
|  | mz_2E-5 | 1953 | 1913 | 40 | 921 | 637 | 636 | 1 | 0 |
|  | mz_5E-5 | 1955 | 1624 | 331 | 926 | 642 | 634 | 8 | 5(3*) |
|  | mz_1E-4 | 1960 | 1098 | 862 | 928 | 646 | 631 | 15 | 11(6*) |
|  | mz_1E-3 | 2083 | 21 | 2062 | 954 | 632 | 522 | 110 | 587(107*) |
| Met_15 | Blank | 2278 |  |  | 26 | 20 |  |  |  |
|  | mz_1E-5 | 2278 | 2278 | 0 | 26 | 20 | 20 | 0 | 0 |
|  | mz_2E-5 | 2278 | 2278 | 0 | 26 | 20 | 20 | 0 | 0 |
|  | mz_5E-5 | 2277 | 2202 | 75 | 26 | 20 | 20 | 0 | 0 |
|  | mz_1E-4 | 2279 | 1724 | 555 | 25 | 19 | 19 | 0 | 0 |
|  | mz_1E-3 | 2329 | 2 | 2327 | 31 | 21 | 19 | 2 | 13(2*) |
| Met_16 | Blank | 75 |  |  | 9 | 8 |  |  |  |
|  | mz_1E-5 | 75 | 75 | 0 | 9 | 8 | 8 | 0 | 0 |
|  | mz_2E-5 | 75 | 75 | 0 | 9 | 8 | 8 | 0 | 0 |
|  | mz_5E-5 | 75 | 73 | 2 | 9 | 8 | 8 | 0 | 0 |
|  | mz_1E-4 | 75 | 60 | 15 | 9 | 8 | 8 | 0 | 0 |
|  | mz_1E-3 | 71 | 1 | 70 | 9 | 8 | 8 | 0 | 3 |

* means the number of unique compounds annotated by unique features

**Fig. s5.** **The summary of intensities fold-change (FC) of consensus features in all groups of files with truncated *m/z* data.** The box chart shows *Log_2_* transformed FC values in all groups (2x10^-4^, 2x10^-3^, 8x10^-3^, 2x10^-2^, 2x10^-1^).

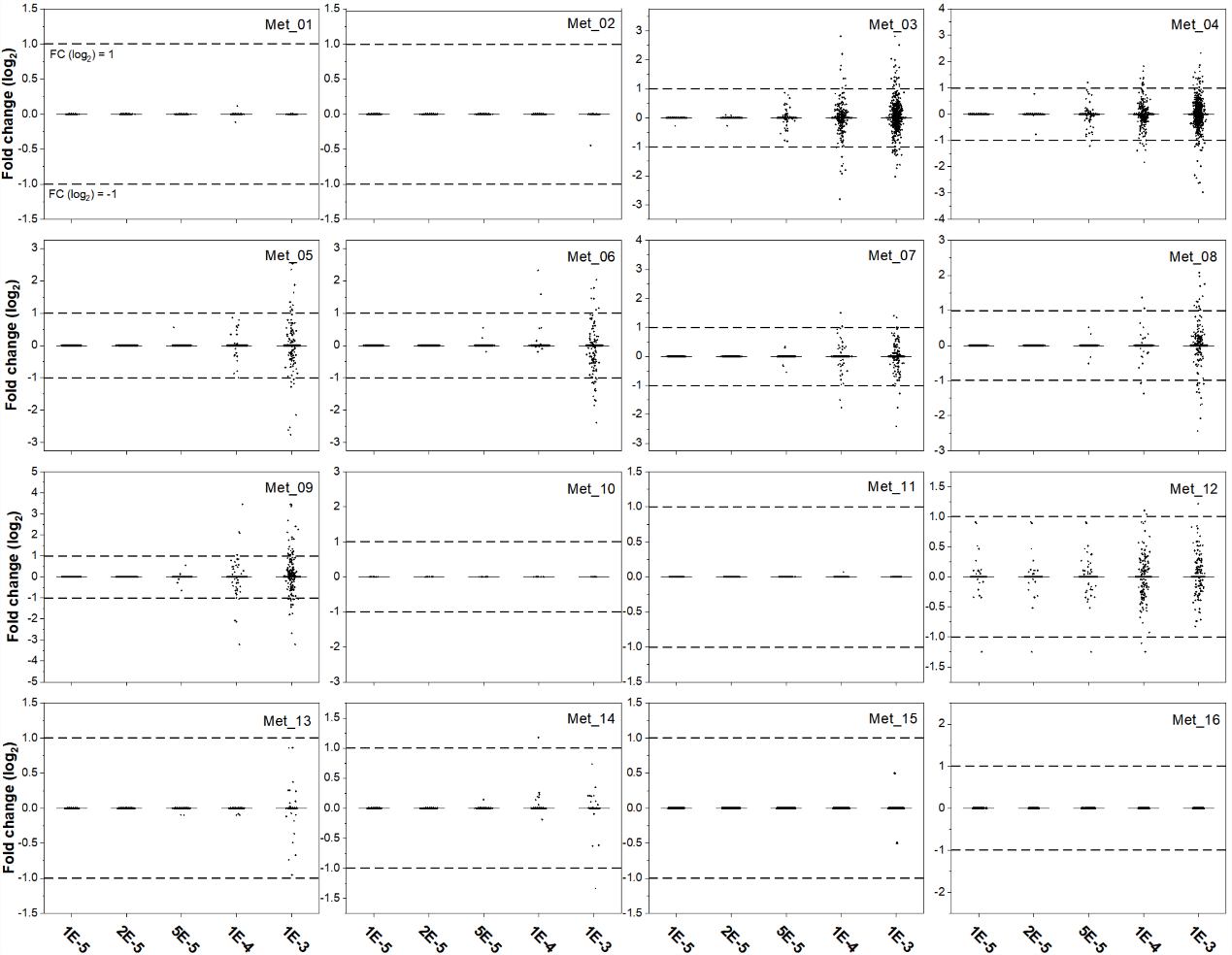

We further examined the intensity FC distribution of consensus features in all groups of files, and these significant features with FC > 2 or < 0.5 were outliers above the line (log2 value equals 1) or below the line (log2 value equals -1) (Figure10). For significant features, a small fraction of less than 0.5% occurs in all groups, except about 0.65% (Met_14) in the group of 10-2. The less significant features were located between the two lines and represented a similar less than 5% of the number of detected features in all groups.

**References**

1.	Alka, O., et al., *DIAMetAlyzer allows automated false-discovery rate-controlled analysis for data-independent acquisition in metabolomics.* Nat Commun, 2022. **13**(1): p. 1347.

2.	Li, Z., et al., *Comprehensive evaluation of untargeted metabolomics data processing software in feature detection, quantification and discriminating marker selection.* Anal Chim Acta, 2018. **1029**: p. 50-57.

3.	John Peter, A.T., M. Peter, and B. Kornmann, *Interorganelle lipid flux revealed by enzymatic mass tagging <em>in vivo</em>.* 2021: p. 2021.08.27.457935.

4.	Suvarna, K., et al., *A Multi-omics Longitudinal Study Reveals Alteration of the Leukocyte Activation Pathway in COVID-19 Patients.* J Proteome Res, 2021. **20**(10): p. 4667-4680.

5.	Yan, Y., et al., *Plasma lipidomics analysis reveals altered lipids signature in patients with osteonecrosis of the femoral head.* Metabolomics, 2022. **18**(2): p. 14.

6.	Tada, I., et al., *Correlation-based deconvolution (CorrDec) to generate high-quality MS2 spectra from data-independent acquisition in multisample studies.* 2020. **92**(16): p. 11310-11317.

7.	Flasch, M., et al., *Stable Isotope-Assisted Metabolomics for Deciphering Xenobiotic Metabolism in Mammalian Cell Culture.* ACS Chem Biol, 2020. **15**(4): p. 970-981.

8.	Muhsen Ali, A., et al., *Metabolomic Profiling of Submaximal Exercise at a Standardised Relative Intensity in Healthy Adults.* 2016. **6**(1): p. 9.

9.	Danne-Rasche, N., S. Rubenzucker, and R. Ahrends, *Uncovering the complexity of the yeast lipidome by means of nLC/NSI-MS/MS.* Anal Chim Acta, 2020. **1140**: p. 199-209.
